# The interplay of additivity, dominance, and epistasis in a diploid yeast cross

**DOI:** 10.1101/2021.07.20.453124

**Authors:** Takeshi Matsui, Martin N. Mullis, Kevin Roy, Joseph J. Hale, Rachel Schell, Sasha F. Levy, Ian M. Ehrenreich

**Author notes:** Correspondence to: Ian M. Ehrenreich, Sasha F. Levy. Equal contribution authors (Takeshi Matsui, Martin N. Mullis).

## Abstract

We used a double barcoding system to generate and phenotype a panel of ~200,000 diploid yeast segregants that can be partitioned into hundreds of interrelated families. This experimental design enabled the detection of thousands of genetic interactions and many loci whose effects vary across families. Traits were largely specified by a small number of hub loci with major additive and dominance effects, and pervasive epistasis. Genetic background commonly influenced both the additive and dominance effects of loci, with multiple modifiers typically involved. The most prominent dominance modifier was the mating locus, which had no effect on its own. Our findings show that the interplay between additivity, dominance, and epistasis underlies a complex genotype-to-phenotype map in diploids.

**One sentence summary:** In diploids, epistasis frequently modifies both additivity and dominance.

---

Most complex traits, including many phenotypes of agricultural, clinical, and evolutionary significance, are specified by multiple loci (*1*). How alleles at these loci collectively produce the heritable trait variation in genetically diverse populations remains unresolved (*2*, *3*). While additive loci play a major role in most traits, non-additive genetic effects are also likely important (*4*–*9*). However, loci with non-additive genetic effects are often difficult to detect, limiting knowledge of their properties (*10*, *11*).

The two purely genetic sources of non-additivity are dominance among alleles of individual loci and epistasis between alleles at different loci (or genetic interactions) (*12*, *13*). Most empirical studies of non-additive genetic effects have focused on haploid or inbred individuals (*14*–*16*), which provide higher statistical power to detect loci due to their nominal levels of heterozygosity. However, by design, these populations cannot furnish insight into dominance or its relationship with epistasis. This is a problem because many eukaryotic species that matter to humans, including our species itself, exist predominantly as diploids that outbreed and have high levels of heterozygosity (*17*–*19*). Dominance may be an important contributor to traits in these species.

When epistasis occurs in diploids, a locus may influence only the additive effects, only the dominance effects, or both the additive and dominance effects of its interactor(s) (*20*–*22*). Such interplay has implications for efforts to genetically dissect phenotypes, predict heritable traits from genotypes, and understand the evolutionary trajectories of beneficial and deleterious alleles. Yet, exploration of the relationship between additivity, dominance, and epistasis has mainly been limited to theory because of technical challenges in identifying non-additive loci.

The budding yeast *Saccharomyces cerevisiae* is a potentially powerful system for studying non-additive genetics in diploids. Haploid yeast segregants with known genotypes can be mated to produce diploids that also have known genotypes (*23*). This strategy facilitates the generation of diploid mapping populations that are roughly the square of the number of haploid progenitors. However, phenotyping large diploid populations of more than ~10,000 individuals has been technically difficult (*23*, *24*), limiting the use of this strategy.

## Phenotyping of a large diploid cross by barcode sequencing

To enable phenotyping of large yeast diploid mapping populations, we developed a system that fuses two genomic barcodes, one from each haploid parent, *in vivo* to create a unique double barcode for each diploid genotype (**Fig. 1A** and **Fig. S1**). We started with two *S. cerevisiae* isolates, the commonly used lab strain BY4716 (BY) and a haploid derivative of the clinical isolate 322134S (3S). These strains differ at ~45,000 SNPs (~0.4% of genome) (*25*, *26*). To ensure segregation of the mating locus, both BY *MAT**a*** × 3S *MAT*α and 3S *MAT**a*** × BY *MAT*α crosses were performed using isogenic strains that had been mating type switched. From these crosses, 600 *MAT*α and 400 *MAT**a*** segregants from distinct four-spore tetrads were marked at the neutral *YBR209W* locus by integrating a random barcode (*27*, *28*). At least two uniquely barcoded strains were recovered per haploid segregant and the genome of each segregant was sequenced to define the genotype represented by each barcode (**Fig. S1** and **Fig. S2A**).

**Fig. 1.**
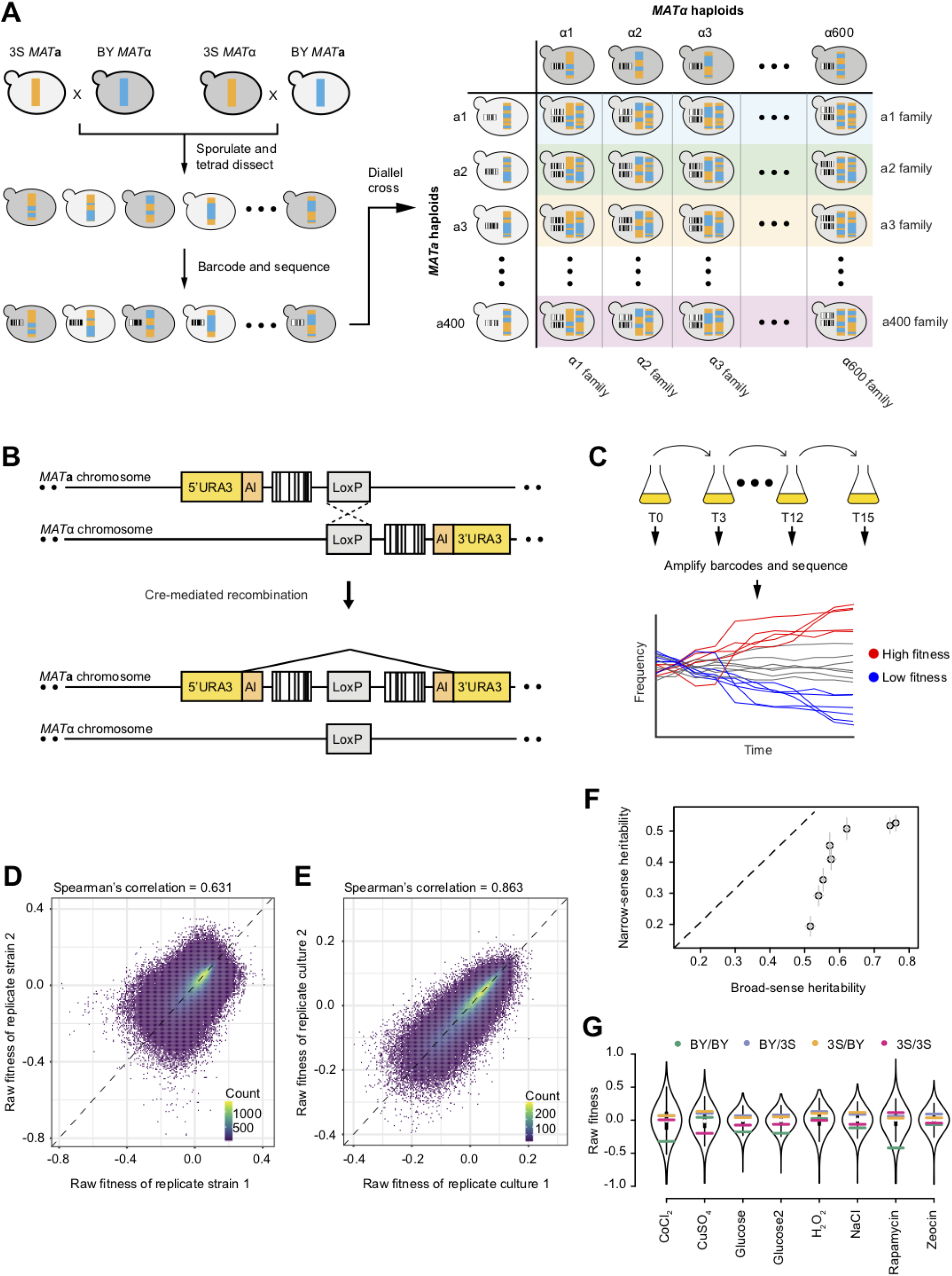
Generating a large panel of diploid segregants with known genotypes that can be phenotyped as a pool. **A.** Overview of the experimental design. Parental haploids, BY and 3S, were mated and sporulated. The resulting MATα and MATa segregants were barcoded at a common genomic location and sequenced. Segregants were mated as pairs to generate a panel of ~240,000 double-barcoded diploid segregants with known genotypes. All diploid segregants originating from a single haploid parent are referred to as ‘family’. **B.** MATα and MATa barcodes were brought to the same genomic location by inducing recombination between homologous chromosomes via Cre-loxP. **C.** Diploid segregants were pooled and grown in competition for 12-15 generations. Barcode sequencing over the course of the competition was used to estimate the fitness of each strain. **D.** Density plot of the raw fitness of double barcodes representing the same genotype in the same pooled growth condition (Glucose 1). **E.** Density plot of the mean raw fitness of the same genotype measured in two replicate growth cultures (Glucose 1 and Glucose 2). **F.** The broad-sense and narrow-sense heritability estimates for the 8 environments. The standard errors for both heritability estimates are shown as error bars for each point. **G.** Violin plots of the fitnesses of diploid segregants in 8 environments. Raw fitness estimates of BY/BY, BY/3S, 3S/BY, and 3S/3S diploid segregants are shown as colored lines.

*MAT**a*** strains and *MAT*α strains (2 barcodes per segregant) were mated as pairs and grown on media that induced site-directed recombination between the *MAT**a*** barcode and *MAT*α barcode on homologous chromosomes (**Fig. 1B** and **Fig. S1E**) (*28*, *29*). This process resulted in a double barcode on one chromosome that uniquely identifies both parents of a diploid segregant and therefore its presumptive genotype (**Fig. S2B**). Using similar methods, we also constructed BY/BY, BY/3S, 3S/BY, and 3S/3S parental diploids.

After the matings, diploids were pooled and competed in seven conditions: cobalt chloride, copper sulfate, glucose, hydrogen peroxide, sodium chloride, rapamycin, and zeocin with the glucose condition performed twice (**Table S1**). Cells were grown for ~15 generations in serial batch culture, with 1:8 dilution every ~3 generations and a bottleneck population size greater than 2 × 10^9^ cells (**Fig. 1C**). Double barcodes were enumerated over 4-5 timepoints by sequencing amplicons from the double barcode locus, and the resulting frequency trajectories were used to estimate the relative fitness of each strain (*30*, *31*).

We recovered on average 197,267 strains per environment with a minimum of two replicate fitness estimates (**Table S2**). Fitness measures correlated well between different barcodes marking the same strain within a growth pool (0.524 < *r* < 0.8 Spearman’s correlation, **Fig. 1D** and **Fig. S3-4**), as did fitness measures of the same barcoded strain assayed in replicate glucose growth cultures (Spearman’s correlation = 0.863, **Fig. 1E**).

Substantial phenotypic diversity was observed in every environment. The majority of this variation was due to genetic factors: broad-sense heritabilities were on average 61% (52-76% across environments), with 40% (19-53%) being additive and 21% (19-26%) being non-additive (**Fig. 1F, Fig. S5,** and **Table S3**). Every environment contained many diploids with more extreme fitness than either the BY/BY or 3S/3S parent (i.e., transgressive segregation). BY/3S and 3S/BY segregants were more fit than the BY/BY or 3S/3S diploids in all environments but one (i.e., heterozygote advantage, **Fig. 1G**).

### Mapping within interrelated families increases statistical power

Using quantile normalized fitness estimates from barcode sequencing, we mapped loci that contribute to growth (**Fig. S6**). Due to our experimental design, diploids generated from the same haploid parent (families) are more genetically related than diploids generated from different parents (**Fig. 1A**). Such family structure causes false positives in genetic mapping. Here, we found that most sites throughout the genome exceeded nominal significance thresholds when fixed effects linear models were applied in a given environment (**Fig. S7**). To enable mapping despite the family structure, we used mixed effects linear models, which are commonly employed in genetic mapping studies involving populations in which individuals show nonrandom relatedness. Specifically, we used Factored Spectrally Transformed Linear Mixed Models (FaST-LMM) (*32*, *33*) to identify an average of 17 loci per environment (10 to 26).

Contrary to expectations that larger sample sizes should yield better statistical power, and therefore, more detections, the numbers of loci identified here were comparable to studies that were at least 60-fold smaller (*2*, *14*, *23*). There are multiple potential, non-mutually exclusive explanations for this observation. For example, statistical controls for family structure may result in a high rate of false negatives or the effects of loci may depend on genetic background. To bypass these difficult to disentangle issues, each of which may impact statistical power, we performed linkage mapping using an alternative strategy that did not require explicitly controlling for family structure. Fixed effects linear models were conducted individually within each of 392 families of diploids that descended from distinct *MAT**a*** parents and consisted of ~600 individuals each.

The family-level scans yielded an average of 16.3 detections per family (**Fig. 2A-B** and **Fig. S8**), which were largely reproducible across replicate cultures (**Fig. 2C**). Detections across the families were then consolidated using 95% confidence intervals, resulting in approximately 58.3 distinct loci per environment (49 to 65), >2.5-fold more loci than detected by FaST-LMM (**Fig. 2A, B,** and **D**). These loci included ~85% of the loci detected using FaST-LMM, implying that the results of family tests encompass those of conventional approaches using aggregate data. Loci identified only in family tests had an average effect that was around one-third of loci detected by both FaST-LMM and family tests (0.08 vs. 0.25; **Fig 2E**). This implies that despite smaller sample sizes, mapping within families provided greater statistical power than mapping in the aggregate data while controlling for relatedness.

**Fig. 2.**
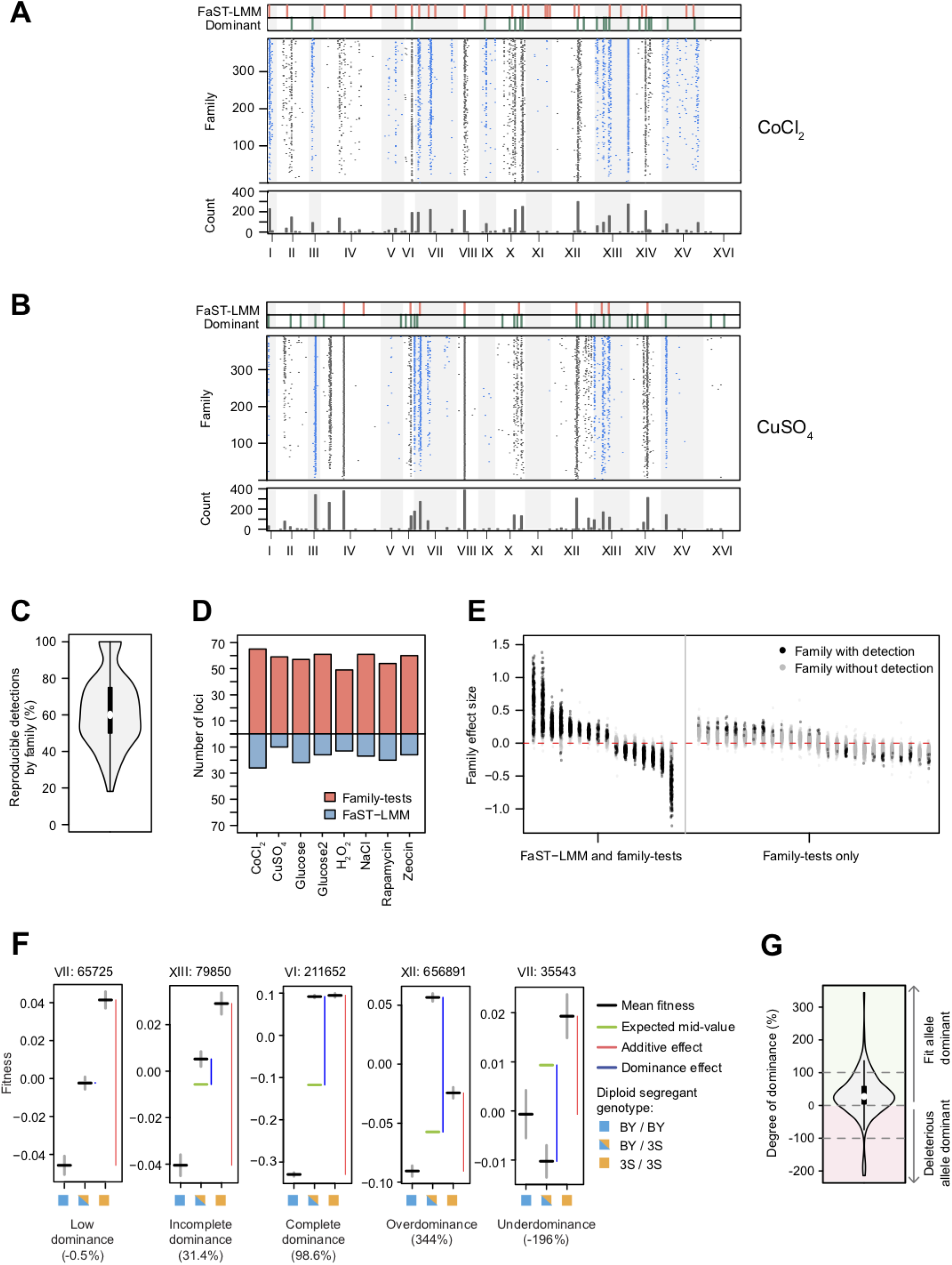
Identification of loci that affect fitness. **A-B.** Loci mapped in CoCl_2_ (**A**) and CuSO_4_ (**B**). Panels from top to bottom are 1) loci detected using the mixed effects linear model FaST-LMM (red bars), 2) loci with dominance effects detected using a fixed effects linear model on the non-additive portion of each diploid’s phenotype (green bars), 3) loci detected using family-tests (black or blue points), and 4) the total number of detections across families for each 50 kb interval (grey bars). **C.** Violin plot showing the % of loci that were detected in both glucose replicates for each family. **D.** Barplot of the number of loci detected by family-tests (red) and FaST-LMM (blue). **E.** Family-level effect size (3S/3S – Het or Het – BY/BY) of loci detected with FaST-LMM in CoCl_2_ (left panel) and family-tests (right panel). Colors indicate whether an effect was detected (black) or undetected (gray) in a family-level scan. **F.** Examples of loci with only additive effect (or low dominance), incomplete dominance, complete dominance, overdominance, and underdominance. Black lines are the mean fitness of diploids subsetted by the genotype state at the focal locus. Gray lines are the standard errors. Green lines are the expected mean fitness of heterozygotes assuming no dominance. Genotype state at each locus is denoted by colored boxes: BY/BY (blue), 3S/3S (orange), is BY/3S (half blue, half orange). Dominance and additive effects (blue and red bars, respectively) for each subset of the data are shown next to the relevant genotype classes. The degree of dominance at a locus is included in parentheses. **G**. Violin plot showing the degree of dominance for all loci detected in the dominance scan. Loci with positive values are dominant towards the allele conferring higher fitness (green), while loci with negative values are dominant towards the deleterious allele (red). All loci with degree of dominance >100% or <−100% exhibit overdominance and underdominance, respectively.

### Loci frequently show dominance effects

In diploids, non-additivity can arise due to dominance among alleles at the same locus, epistasis between alleles at different loci, or a mixture of the two. To identify such non-additive loci from the aggregate data, we extracted the non-additive portion of each diploid’s phenotype (**Fig. S9**) (*23*). Using these values accounts for family structure and enables mapping of non-additive loci in the full segregant panel with fixed effects linear models. Regarding dominance effects, we identified an average of 18 loci showing dominance per environment (12 to 30). Only 45% of these loci were also identified by FaST-LMM, while 82% of these loci were detected in the family-level scans. Among the loci with dominance effects, the average degree of dominance was ~51% (i.e., heterozygotes’ fitnesses were roughly halfway between the average of the two homozygotes and one of the homozygotes), with 82% of the loci showing incomplete dominance (**Fig. 2F-G**). Only ~7% of the loci exhibited complete dominance, while overdominance (~8%) and underdominance (~3%) were seen among the remaining loci with dominance effects. ~77% of the loci showed dominance towards the allele conferring higher fitness (**Fig. 2G**), which may explain why segregants were more fit than the BY/BY or 3S/3S diploids(**Fig. 1G**).

### Epistatic hubs govern both additivity and non-additivity

We also used the non-additive portion of phenotype to perform comprehensive genome-wide scans for genetic interactions. We identified an average of 440 two-locus interactions per environment (377 to 538) (**Fig. 3A** and **Fig. S10**). Our large sample size had a pronounced impact on detection: ~40-fold more interactions per environment were detected than previous studies that phenotyped smaller mapping populations using conventional approaches (*14*, *23*). Our large sample size also enabled comprehensive scans for three-locus interactions with a reduced set of markers, identifying an average of 6,152 per environment (4,845 to 7,301) (**Fig. 3A** and **Fig. S11**). Loci involved in three-locus interactions were identified across all chromosomes and distributed widely throughout the genome.

**Fig. 3.**
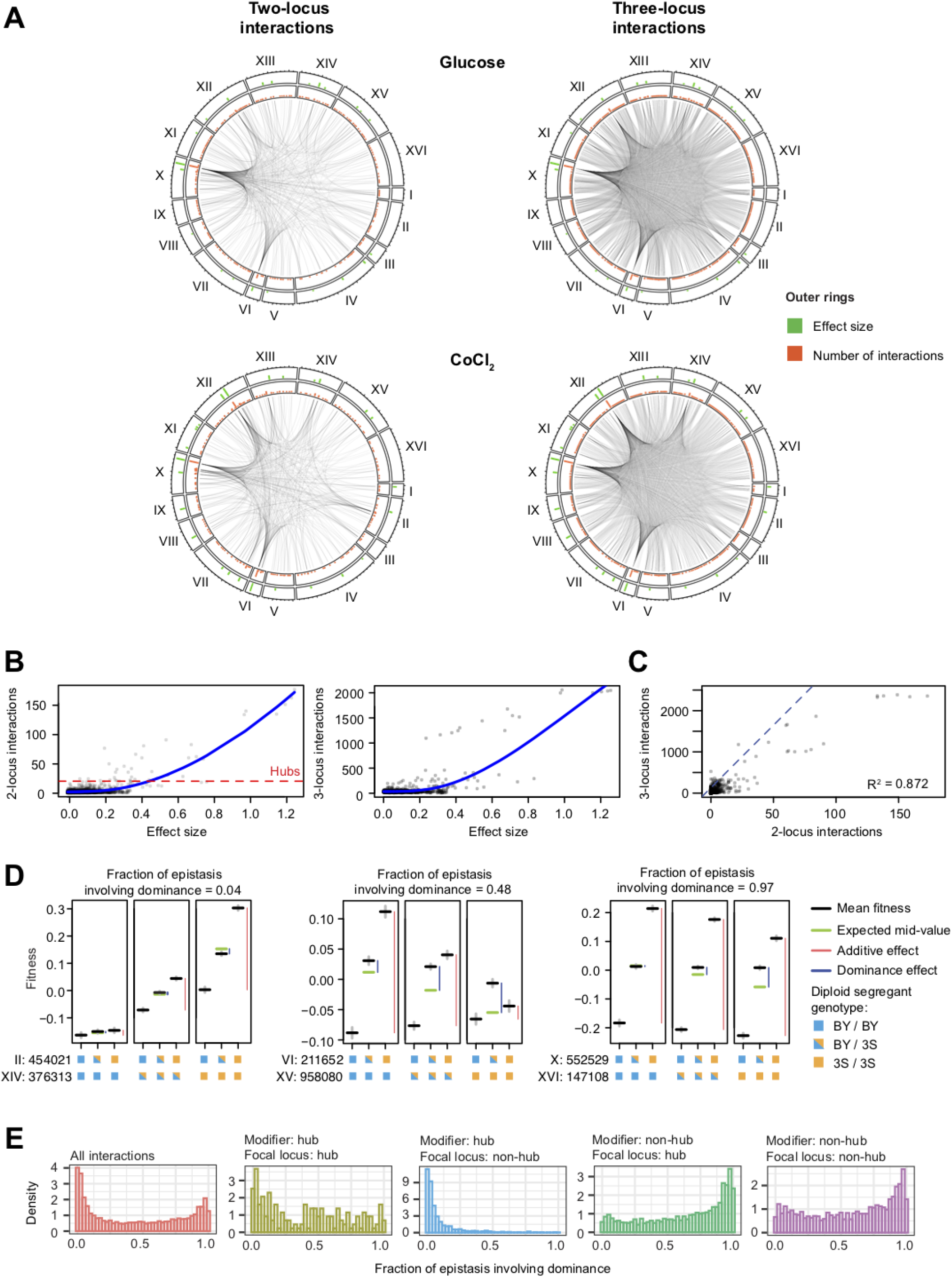
Interactions often affect both the additive and dominance effects of involved loci. **A.** Interaction plots of all two-locus (left) and three locus (right) effects for two representative environments. Significant interactions between loci are shown as connecting lines. Green bars are the effect size of a locus, calculated as the absolute difference between the mean fitness of diploids that are 3S/3S and BY/BY at the focal locus. Orange bars are the number of interactions detected for each locus. **B.** Scatter plot of the absolute effect size of a locus and the number of two-locus (left) and three-locus (right) interactions in which it is involved. Local regressions are shown as blue lines. **C.** Scatter plot of the number of two-locus and three-locus interactions per locus. **D.** Examples of genetic interactions with different fractions of epistasis involving dominance. Black lines are the mean fitness of diploids subsetted by the genotype state at the two involved loci. Gray lines are the standard errors. Green lines are the expected mean fitness of heterozygotes assuming no dominance. Genotype state at each locus is denoted by colored boxes: BY/BY (blue), 3S/3S (orange), is BY/3S (half blue, half orange). The first locus is the locus whose effect is being modified, and the second locus is the modifier locus. Dominance and additive effects (blue and red bars, respectively) for each subset of the data are shown next to the relevant genotype classes. **E.** Density plot of the fraction of epistasis involving dominance for all interactions (red), hub--hub (yellow), non-hub--hub (blue), hub--non-hub (green), and non-hub--non-hub interactions (purple).

We next analyzed the relationship between individual loci and their genetic interactions. We found a strong positive relationship between the effect of a locus and its involvement in two- and three-locus interactions (**Fig. 3B**). This suggests that loci with larger effects tend to genetically interact with many loci or that their interactions are easier to detect. We also observed a clear linear relationship between the number of two- and three-locus interactions of a given locus (**Fig. 3C**). Notably, certain loci exhibited many more interactions than others, acting as ‘hubs’ (here defined as loci with >20 two-locus interactions in at least one environment) (*8*). On average, ~4.5 hubs were detected per environment, and the same hub was often detected in multiple environments. A majority (>54%) of all two- and three-locus interactions involved at least one hub. Fine-mapping localized the Chromosome VI, VIII, X, and XII hubs to genes involved in amino acid sensing (*PTR3*), copper resistance (*CUP1*), vacuolar protein sorting (*VPS70*), and a gene of unknown function (*YLR257W*), respectively.

### Relationships between epistasis and dominance in diploids

In haploids, epistasis can only influence the additive effect of a locus because there are no heterozygotes. In diploids, however, epistasis can modify a locus’ additive effects, dominance effects, or both additive and dominance effects (*20*–*22*). To better characterize how loci are modified, each two-locus interaction was partitioned into additive and dominance components. We found that changes in dominance account for ~44% of the average epistatic effect (**Fig. 3D-E**), implying that interactions often affect both additivity and dominance. However, this fraction varied depending on whether the modifying locus was a hub. When the modifier was a hub, dominance accounted for little of the epistatic effect (11.9% on average), implying that hubs mostly modify the additive component of the interacting loci. By comparison, when the modifier was not a hub, epistasis was mostly composed of dominance (64% of interactions had a larger dominance component). These data suggest that epistasis commonly involves modification of dominance in diploids and that hubs act in a distinct manner from loci that are not hubs.

We next examined how the additive and dominance effects of hubs were modified by genetic interactions. In most cases, hubs genetically interacted with a small number of major effect modifiers and many minor effect modifiers (**Fig. 4A**). The major effect modifiers typically influenced only the additive or only the dominance effect of a hub, suggesting that distinct sets of loci govern additive and dominance effect sizes (**Fig. 4A**). Whereas the most frequent major effect modifiers of the additive effects of hubs were other hubs (**Fig. 4B**), the single most frequent major effect modifier of the dominance effects of hubs was a locus on Chromosome III. Collectively, multiple modifier loci could cause a hub locus to show a broad range of effect sizes across different genetic backgrounds (**Fig. 4C**).

**Fig. 4.**
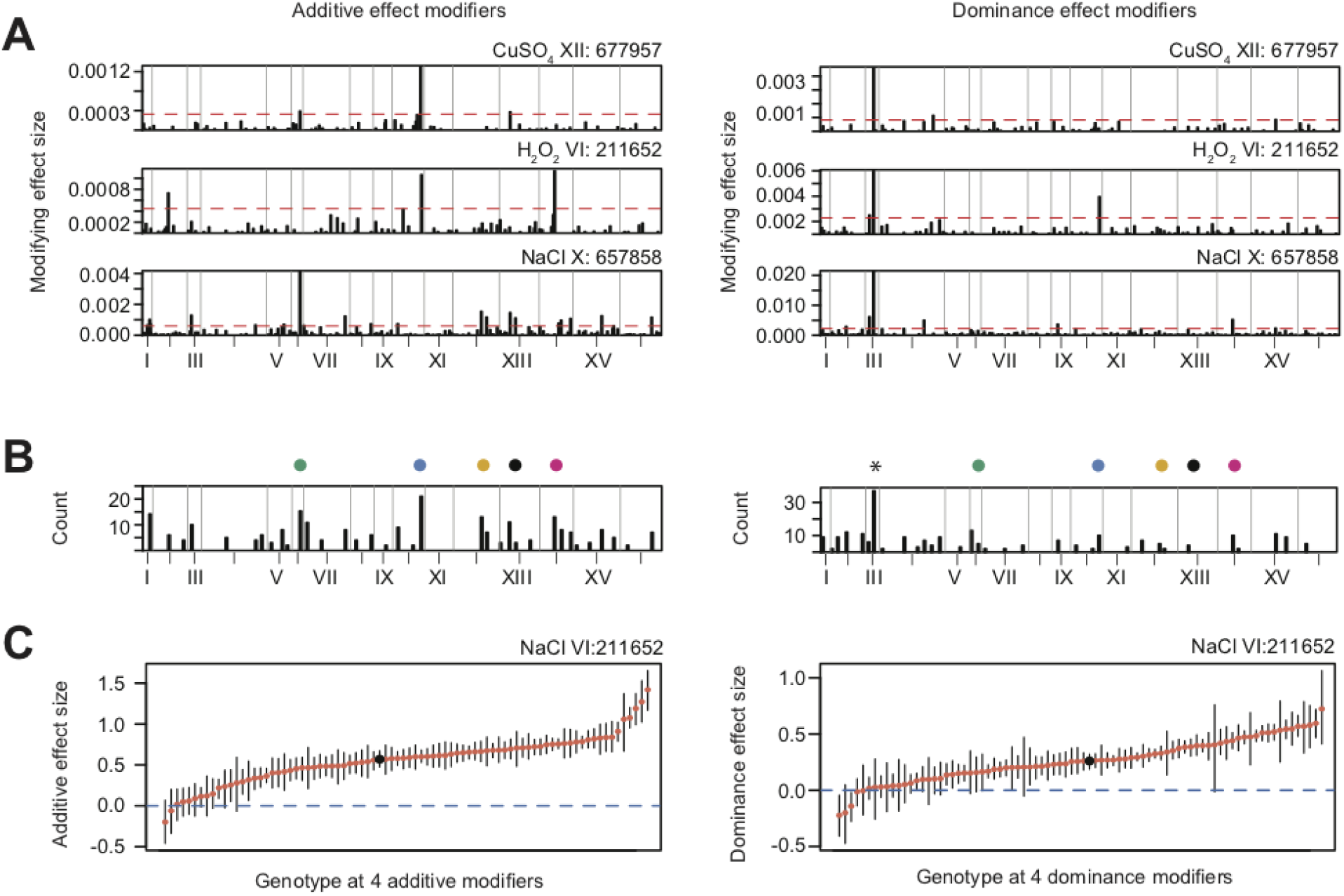
Multiple modifier loci cause hubs to exhibit a range of effect sizes across different genetic backgrounds. **A.** Specific examples of hubs on chromosome VI, X, and XII (each row) and their additive (left) and dominance (right) modifiers. The height of the bar corresponds to the magnitude of the modifying effect. The dotted red line shows the threshold in which loci were considered as major effect modifiers. **B.** Barplot showing the total number of times loci were detected as a major effect modifier of additive (left) or dominance (right) effects of hubs across environments. Colored dots indicate hub loci. An asterisk indicates a non-hub locus on ChrIII. **C.** Additive (left) or dominance (right) effect size of chromosome VI hub across different allelic combinations of its 4 largest effect modifiers. Red points are the effect size of a genotype class based on the genotype state of the 4 modifiers. Black point is the overall effect size of the locus. Black lines are bootstrapped 95% confidence intervals.

### Characteristics of the Chromosome III dominance modifier

Although not a hub, the Chromosome III locus nevertheless had a prominent impact on phenotype by modifying the dominance effects of multiple variable effect loci. Interactions with the Chromosome III locus had greater impacts on dominance than additivity at all focal loci. For example, in hydrogen peroxide, dominance at the Chromosome X variable effect locus depended on the Chromosome III locus, ranging from complete to nearly absent in a genotype-dependent manner (**Fig. 5A**). We delimited the Chromosome III locus to a 3 kb region containing the mating locus and a few other genes (*BUD5*, *TAF2*, and *YCR041W*).

**Fig. 5.**
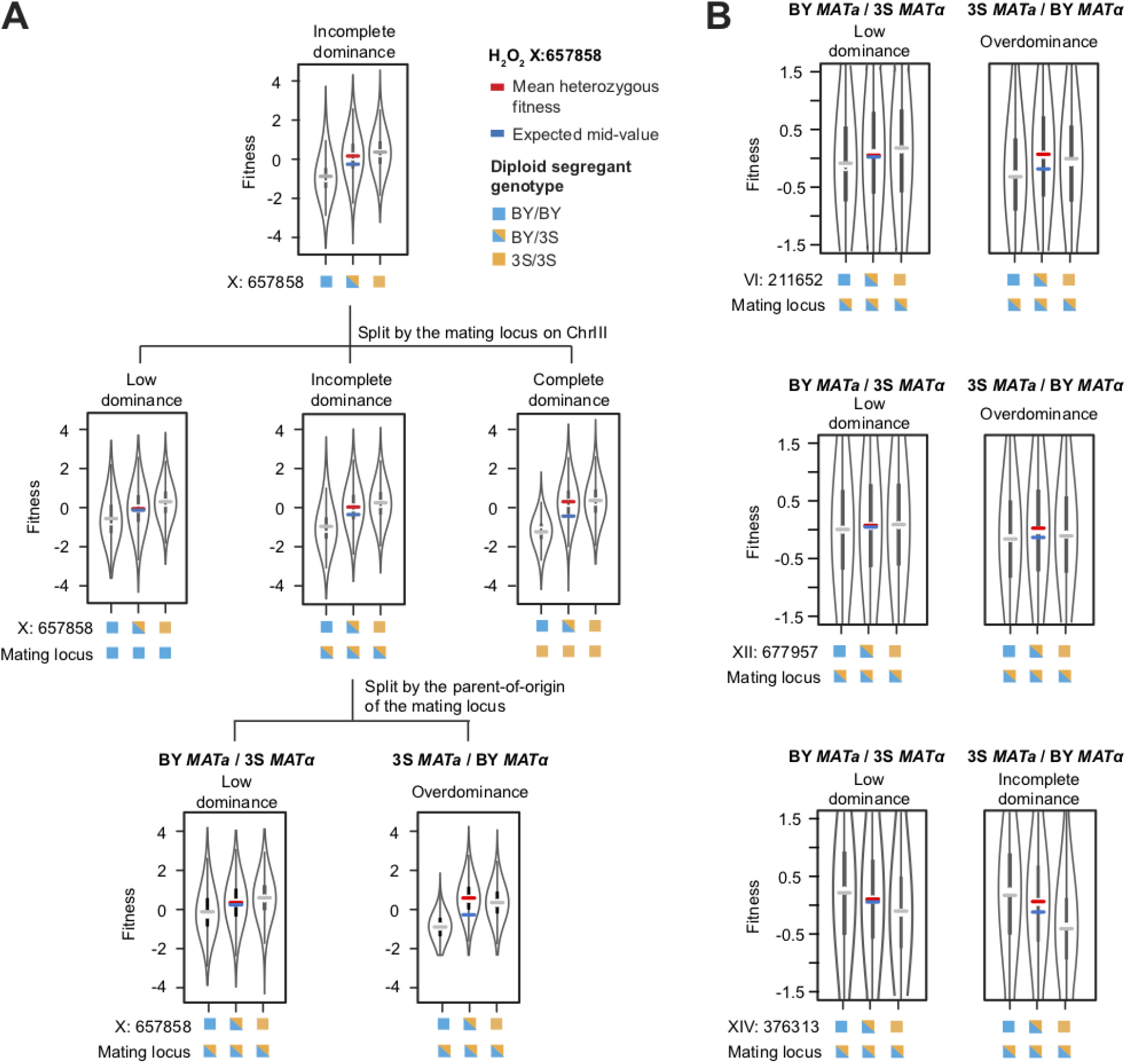
Parent-of-origin of the mating locus influences dominance at hubs. **A.** Violin plots of the fitness distribution of diploids split by the genotype at the Chromosome X locus (top), further split by the genotype at the mating locus on Chromosome III (middle), and the parent-of-origin at the mating locus (bottom). Genotype state at each locus is denoted by colored boxes: BY/BY (blue), 3S/3S (orange), is BY/3S (half blue, half orange). Lines are the observed mean fitness in the homozygous genotype classes (gray), the observed mean fitness in heterozygous genotype classes (red), and the expected heterozygous fitness if there was no dominance (blue). **B.** Violin plots of the fitness distribution of diploids split by the genotype state of a hub locus and the parent-of-origin of the mating locus.

Yeast mating types possess different nonhomologous gene cassettes at the mating locus, which encode distinct transcription factors that are master regulators of the *MAT**a***, *MAT*α, and diploid transcriptional programs (*34*). This region of the genome is unique because four genotype classes segregate (BY *MAT**a,*** 3S *MAT**a**,* 3S *MAT*α, and BY *MAT*α), and as a result, the two heterozygotes are not identical (**Fig. S12**). To test if the mating locus is the dominance modifier, we partitioned Chromosome III heterozygotes based on their parents-of-origin for the *MAT**a*** and *MAT*α cassettes and found a difference: dominance was only visible in the 3S *MAT**a*** / BY *MAT*α genotype class (**Fig. 5A**). Other hub loci modified by Chromosome III showed the same relationship between dominance and the parent-of-origin of the mating loci (**Fig. 5B**). These results suggest that BY and 3S harbor functional differences in one or both mating cassettes.

## Discussion

We used a double barcoding system to generate and phenotype an extremely large panel of diploid yeast segregants that can be partitioned into hundreds of interrelated families. This experimental design enabled the detection of thousands of loci, including at least an order of magnitude more genetic interactions than discovered in previous yeast crosses. Analysis of these epistatic loci identified a modest number of hubs that have large effects, show pervasive epistasis, and control most phenotypic variation across environments, as well as many other loci that genetically interact with these hubs.

Genetic background commonly modified the magnitude of, or completely masked, the effects of the hubs, indicating that the largest effect loci identified in mapping studies are highly sensitive to genetic background. Such non-additive genetic background effects are likely to hamper efforts to predict phenotype from genotype by limiting the extrapolation of effect estimates from one genetic context to others. However, our finding that large effect loci were most impacted by other major effect loci does provide some optimism that characterizing a limited set of interactions may account for a substantial portion of these genetic background effects.

Because our experiments were performed in outbred diploids rather than haploids or inbred diploids, we could detect dominance effects and whether dominance is modified by epistasis. We showed that dominance effects are common and that the magnitude of dominance can strongly depend on the alleles of interacting loci. The potential existence of dominance modifiers has been discussed in theory, but to date, only a single dominance modifier has been found in a plant self-incompatibility locus (*35*, *36*). Our results show that dominance modifiers are prevalent and raise the intriguing possibility that sites with atypical allele dynamics within natural populations, the yeast mating locus here and a self-incompatibility locus in plants, are more likely to harbor dominance modifiers with major effects.

Generally, we found that heritable traits in yeast are more genetically complex than formerly appreciated. Relative to the cross that we examined, natural populations may harbor substantially higher genetic diversity, meaning traits could be even more complex and difficult to dissect. Our work supports the premise that, to the extent possible, focusing on groups of more closely related individuals, such as the families studied here, can enhance statistical power and precision relative to populations with greater diversity (*23*, *24*, *37*). The genetic insights gained from these more closely related groups can then be leveraged to inform the genetic architecture of traits in more diverse populations in which many critical genetic effects may otherwise be obscured.

## Acknowledgments

The authors thank members of the Ehrenreich and Levy labs for feedback on drafts of this manuscript and Oscar Aparicio for an *HO* plasmid.

